# An Open Access Resource for Marmoset Neuroscientific Apparatus

**DOI:** 10.1101/2024.11.12.623252

**Authors:** Isabela Zimmermann Rollin, Daniel Papoti, Mitchell Bishop, Diego Szczupak, Michael R. Corigliano, T. Kevin Hitchens, Bei Zhang, Sarah K. A. Pell, Simeon S. Guretse, Audrey Dureux, Takeshi Murai, Stacey J. Sukoff Rizzo, L. Martyn Klassen, Peter Zeman, Kyle M. Gilbert, Ravi S. Menon, Meng-Kuan Lin, Stefan Everling, Afonso C. Silva, David J. Schaeffer

## Abstract

The use of the common marmoset (*Callithrix jacchus*) for neuroscientific inquiry has grown precipitously over the past two decades. Despite windfalls of grant support from funding initiatives in North America, Europe, and Asia to model human brain diseases in the marmoset, marmoset-specific apparatus are of sparse availability from commercial vendors and thus are often developed and reside within individual laboratories. Through our collective research efforts, we have designed and vetted myriad designs for awake or anesthetized magnetic resonance imaging (MRI), positron emission tomography (PET), computed tomography (CT), as well as focused ultrasound (FUS), electrophysiology, optical imaging, surgery, and behavior in marmosets across the age-span. This resource makes these designs openly available, reducing the burden of de novo development across the marmoset field. The computer-aided-design (CAD) files are publicly available through the Marmoset Brain Connectome (MBC) resource (https://www.marmosetbrainconnectome.org/apparatus/) and include dozens of downloadable CAD assemblies, software and online calculators for marmoset neuroscience. In addition, we make available a variety of vetted touchscreen and task-based fMRI code and stimuli. Here, we highlight the online interface and the development and validation of a few yet unpublished resources: Software to automatically extract the head morphology of a marmoset from a CT and produce a 3D printable helmet for awake neuroimaging, and the design and validation of 8-channel and 14- channel receive arrays for imaging deep structures during anatomical and functional MRI.

## 1. Introduction

Resources to support the use of the common marmoset (*Callithrix jacchus*) for neuroscientific inquiry have developed rapidly ^1–12^, with this small New World primate species poised to inform translational gaps between rodents and humans ^13–16^. One of the many practical advantages of the marmoset model is that preclinical instruments originally designed for rodent models can be ported for use in marmosets, such as small-bore MRI or PET systems, electrodes, and optical imaging setups^17–20^. Despite the small marmoset body (∼350-400 g as adults) notionally fitting within these apparatus, rodent-specific hardware is often incompatible with the architecture of the marmoset musculoskeletal system, which is fundamentally different than that of a rodent^21^. As a semi-arboreal species, marmosets have evolved longer proximal limb bones and a neuraxis orientation more characteristic of anthropoids, with an enlarged cranial cavity to support a larger brain with forward-facing eyes for stereoscopic vision^22,23^. Indeed, one of the key benefits of the marmoset neuroscientific model is its well-developed visual system and concomitant frontoparietal cortical architecture^23–27^ which is amenable to the recapitulation of human brain diseases. To support these efforts, our collective laboratories and others from across the globe have invested significant effort in porting – or more often, designing new – neuroscientific apparatus for the marmoset.

The purpose of this resource (Marmoset Neuroscientific Apparatus; MNSA: https://www.marmosetbrainconnectome.org/apparatus/) is to centralize those efforts under the auspice of the Marmoset Brain Connectome (MBC; marmosetbrainconnectome.org) and the NIA Open Science MARMO-AD consortium. In line with the 3-Rs (replacement, reduction, refinement)^28^, this resource aims to make these detailed designs openly available and editable, reducing the burden of de novo development across the marmoset field and easing financial burdens on funding agencies by minimizing duplicate efforts and promoting shared resources. Centralizing these designs also allows for improved repeatability of studies across labs and institutions, with this level of detail rarely included in published methods.

One area of our collective focus has been developing radiofrequency coils for ultra-high field preclinical MRI systems, offering exquisite signal-to-noise ratio and resolution^1,29–32^. These developments have enabled the generation of structural and functional marmoset brain atlases^3–6,9–12^ and allowed for insights into where marmoset brain structure and function fit with reference to other preclinical modeling species^20,27,33–35^. With clear effects of anesthesia on functional signals in the marmoset^36,37^ and the advent of conducting task-based fMRI in awake-behaving marmosets, there has been prodigious bias toward engineering apparatus to support comfortable – yet motion-free – imaging in the marmoset. To that end, our groups and others have developed MRI hardware to support both surgically implanted and otherwise naïve, noninvasive designs^1,9,29–32,38–40^. More recently, to support high-throughput translational studies through which hundreds of marmosets are imaged, we developed AMIHGOS (Automated Marmoset Imaging Helmet Generator, an Open-Source Software) to automatically generate anatomically conformed 3D printable helmets based on computerized tomography (CT) images of individual animals. This software allows for generation of a comfortable animal-specific helmet that can be used for fully awake MRI, PET, CT, and likely other modalities that we have yet to fully explore. AMIHGOS supports the fitting of a helmet for the sphinx position (the natural resting position of a marmoset) that is optimized to attach to a ‘multimodal’ marmoset holder that minimizes the experimenters’ handling of the marmoset. Together, this apparatus fits within small-bore MRI, PET, and CT systems and can be coupled with a conformal radiofrequency receive array. Here, we detail the development of the AMIHGOS software, designed to maximize comfort during awake scanning while also minimizing motion. Further, we detail the design and testing of two radiofrequency coils, one for fully awake marmoset fMRI (8 channels) and the other optimized for anatomical scanning in stereotactic position under inhalant anesthesia (14 channels).

An ongoing field of inquiry in marmoset neuroscience is exploring the extent and limits of their cognitive abilities, including aspects such as memory capacity, problem-solving skills, and the presence of advanced social cognition. To that end, we have designed a variety of tasks that can be deployed and assayed using touchscreen systems^41–43^ or during awake behaving fMRI^35,44^. Further, we have developed ‘multi-marmoset’ systems to allow for simultaneous functional MRI^45^ or PET to disentangle the neural correlates of dyadic social interactions. Such complex behaviors are often better studied with techniques biased in the time domain using electrophysiology, optical imaging, or even a combination of multiple modalities (e.g., MRI coils that mount directly to surgically implanted head chambers). The circuitries controlling these complex behaviors are further complicated by multisensory integration (e.g., olfaction of pheromones, visual input, auditory input). All such designs are represented through the online platform, from olfactometers, to mirror systems for displaying stimuli and recording eye movements, and coil systems designed for auditory stimulation. Further, the task designs – down to the code and actual stimuli – are available for use, many of which were implemented to acquire published data. Further, marmoset-specific calculators (e.g., for medication calculations) are included not only for easy access, but also to promote standards across the field.

The Marmoset Neuroscientific Apparatus (MNSA) resource aims to serve as a comprehensive platform and repository for advancing marmoset neuroscience research, designed with scalability to adapt alongside scientific progress. The MNSA resource makes available engineering designs, software, data, and tasks that have supported over a decade of marmoset research across multiple institutions. With a dedicated ‘submissions’ page, this resource also enables the sharing and integration of designs contributed by researchers from across the community, fostering collaboration and innovation in marmoset research. More broadly, this resource lowers the barrier for entry for scientists working with other species to transition more readily into marmoset neuroscience and to extend marmoset research beyond imaging and behavioral techniques.

## 2. Materials and Methods

### 2.1. Animals

All procedures in this study were approved by the Institutional Animal Care and Use Committee (IACUC) of the University of Pittsburgh. The marmosets (*Callithrix jacchus*) used in this study were socially housed with two animals per enclosure, provided an ad libitum diet, unrestricted access to water, and enriched environments, including toys and other stimuli. Forty-nine adult marmosets (10 females, 39 males; aged 4.5-138 months, weight 170-552 g) contributed to the data shown in the results, but many others participated in the development of the apparatus across institutions. Unless otherwise specified, experiments shown in this manuscript were conducted with a 9.4 Tesla MRI (9.4T/30cm USR magnet; Bruker-Biospin GmbH, Ettlingen, Germany) or PET/CT (Si78; Bruker BioSpin GmbH, Ettlingen, Germany) system, both at the University of Pittsburgh. If the imaging was conducted in awake marmosets, all animals underwent acclimatization training as described by Silva et al.^46^.

### 2.2 Marmoset Neuroscientific Apparatus Online Interface

The marmoset neuroscientific apparatus online repository is available at https://www.marmosetbrainconnectome.org/apparatus/ (Figure 1A) and houses dozens of designs organized in the following categories: MRI, histology, electrophysiology, behavior, husbandry, optical imaging, CT, PET, surgical apparatus, tFUS, and software. Each design is accompanied by its own subpage (Figure 1B), which includes relevant details on building or otherwise using the apparatus in marmoset neuroscience. In the case of engineering designs, users can view photos, 3D renderings, and rotatable 3D models (Figure 1B), as well as download assembled or individual files in open formats. All designs include stereolithography files (STL), which can readily be 3D printed onsite or outsourced for manufacturing. For those inclined to modify the designs, two neutral CAD design formats are included: Initial Graphics Exchange Specification (IGES files) and Standard for the Exchange of Product Data (STEP files) – with these, users can open and modify assemblies and individual parts in openly available CAD design software, or convert the files for manufacturing (e.g., Computer Numerical Control (CNC) milling). All included designs are actively used in our laboratories and undergo continuous refinement. If a design was used in a scientific publication (even if it was not directly published in the article), the relevant publication is listed on the subpage for proper attribution. Under the “Details” tab, users will find a short description of the design, the reference publication, and contact information for the person(s) responsible for the design. All designs are provided under the Creative Commons BY-NC-SA 4.0 license, such that they can be shared and adapted by others as long as proper attribution is given, they are not used for commercial purposes, and any derivative works are distributed under the same license.

**Figure 1:**
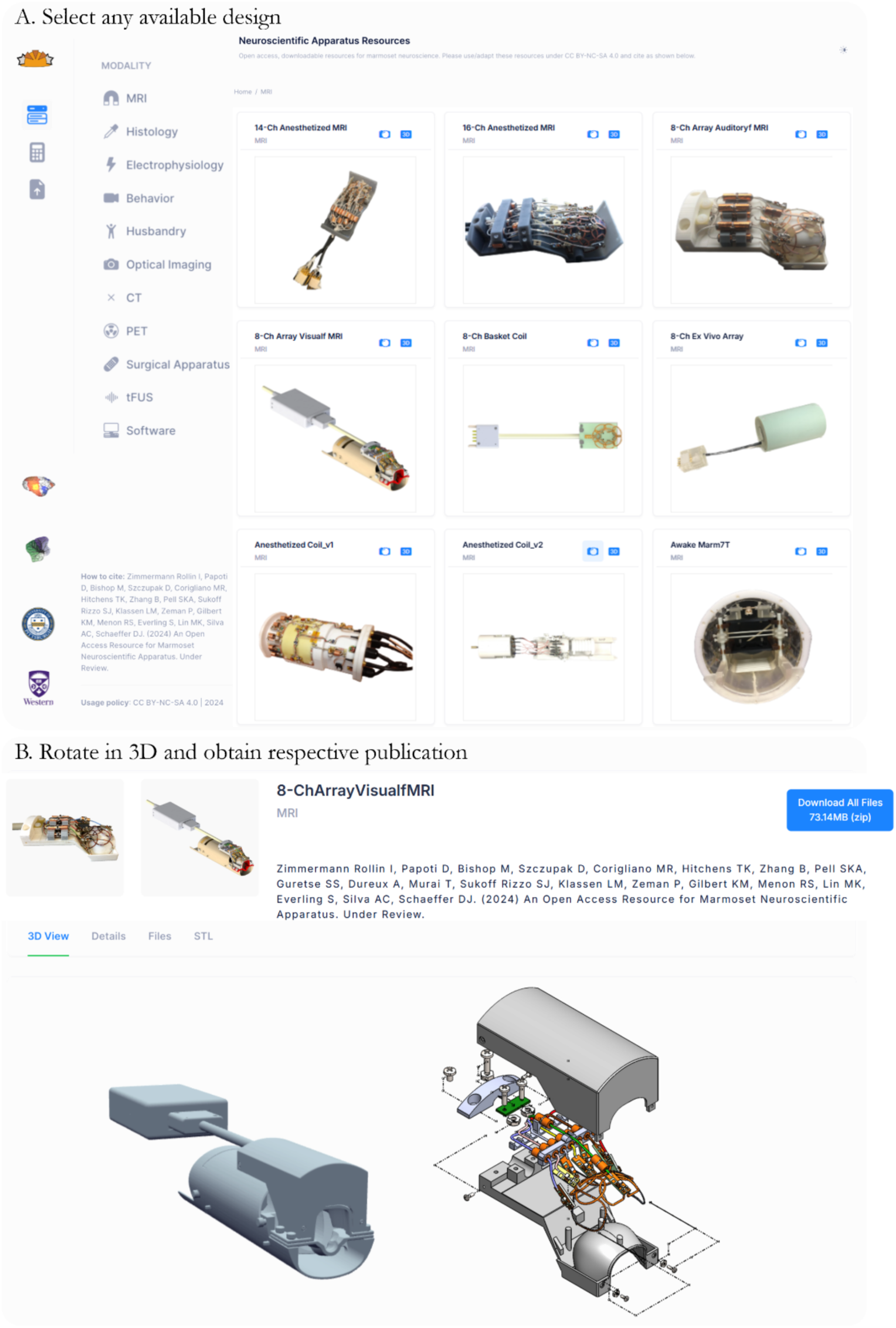
(A) A screenshot of the home page of the Neuroscientific Apparatus Resource. (B) After selecting a design, the user is directed to a page where they can access a zoomable and rotatable 3D rendering along with photos and relevant design files.

In addition to the engineering designs, there are currently three categories that house resources that do not fall under ‘apparatus’ per se. The first, included in the ‘CT’ category, are CT images (Figure 2A) available in raw 3D Neuroimaging Informatics Technology Initiative (NIfTI) and 3D printable formats (extracted skulls in STL format) across the marmoset age span – among other possible uses, one goal of providing these files is for presurgical planning on a 3D printed skull (or digitally) for scientists without ready access to a CT of their animal. By using a provided file from an animal of a similar age and sex, one can estimate surgical trajectories and even practice the surgery in a stereotactic apparatus^47,48^. The second category is ‘software,’ which currently includes paradigms for functional MRI and touchscreen tasks, complete with the code and stimuli (e.g., photos or videos) needed to present the task to a marmoset. While we use a commercial hardware system that was specifically selected so all investigators can purchase the same setup, we also share our task-specific code and ancillary hardware (such as touchscreen nestboxes). Yet another feature of the online resource is the ‘calculator’ tab (Figure 2B), which allows for calculations used across imaging and surgical operations, with some also plotting ranges of values. For example, the ‘medication’ calculator uses weight as an input to instantly compute presurgical, postsurgical, and emergency doses of drugs. This calculator and others were implemented for practical reasons (e.g., accessing on a computer in the experiment room), but also can serve to set standard practice across the field.

**Figure 2:**
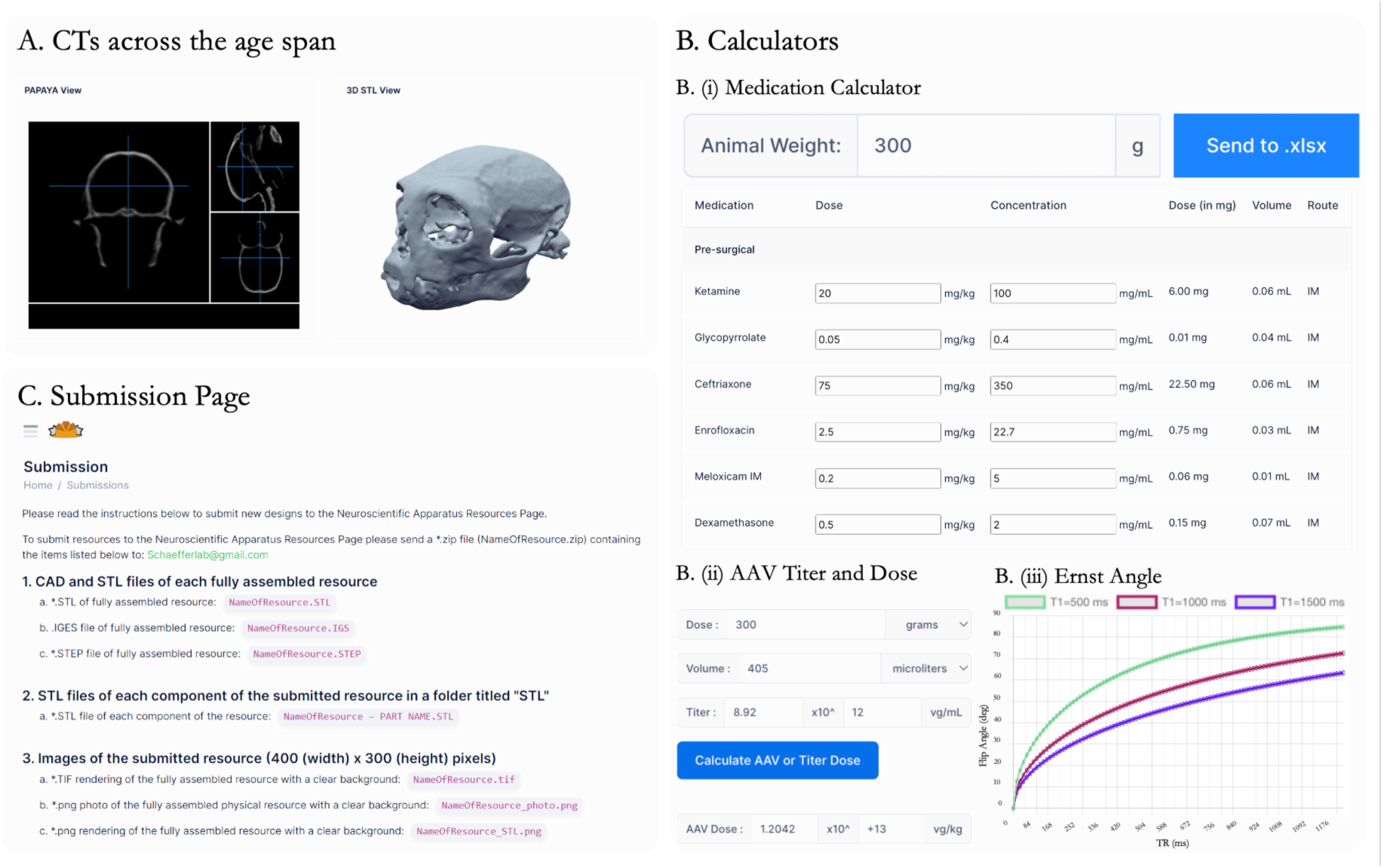
(A) CT images available in an online viewer (also downloadable) across the age span for marmosets of different sexes and weights. (B) Examples of calculators available on the website, including a (i) medication calculator for doses of different agents from a single input (weight of animal), (ii) adeno-associated viral (AAV) injection dose calculator (ii), and (iii) Ernst Angle calculator for optimizing contrast with MR images. (C) Submission Page for groups interested in contributing to the resource.

Researchers interested in submitting a mechanical design or software for inclusion on the website can visit the submission page (Figure 2C), which provides detailed instructions on the required file types and submission process.

### 2.3 Automated Marmoset Imaging Helmet Generator, an Open-Source Software (AMIHGOS)

To support high-throughput translational studies through which hundreds of marmosets are imaged, we developed AMIHGOS (Automated Marmoset Imaging Helmet Generator, an Open-Source Software) to automatically generate anatomically conformed 3D printable helmets based on computerized tomography (CT) images of individual animals. These individualized helmets can be 3D printed using MRI-compatible materials (e.g., Durable Resin; Form 3, Formlabs, Somerville, Massachusetts, USA). After manufacturing, the helmets are lined with a thin layer (1 mm) of ultra-high density foam (Ultra High Density Craft Foam 85 kg/m^3^; The Foamory, Raleigh, NC, USA) to maximize comfort in awake MRI, PET, or CT. Anecdotally, we have learned that marmosets are more likely to move if they are uncomfortable due to a peak or sharp edge in the helmet. As such, the AMIHGOS software is the culmination of years of iterating designs to maximize comfort during long (2+ hour) imaging sessions. This software is continuously improved as we encounter outliers (excessively small or large heads). In short, the software works by thresholding Hounsfield Scale (HU) values from the CT to extract the soft tissue (skin, which is contoured by underlying muscle and bone) and then subtract that shape from a standard helmet template in an optimized position for the sphinx posture.

#### 2.3.1 AMIHGOS processing pipeline

All necessary files and installation instructions for the AMIHGOS software are available under the software category. An overview of the processing steps is illustrated in Figure 3. Once the user has configured the local Python environment, they can launch AMIHGOS from the terminal. The home page allows users to select NIFTI or Digital Imaging and Communications in Medicine (DICOM) formatted CT images as inputs from the file explorer. If the user has already segmented and extracted a mesh of the subject head, they can skip directly to the head mesh transformation and subtraction step by clicking the button below it.

**Figure 3:**
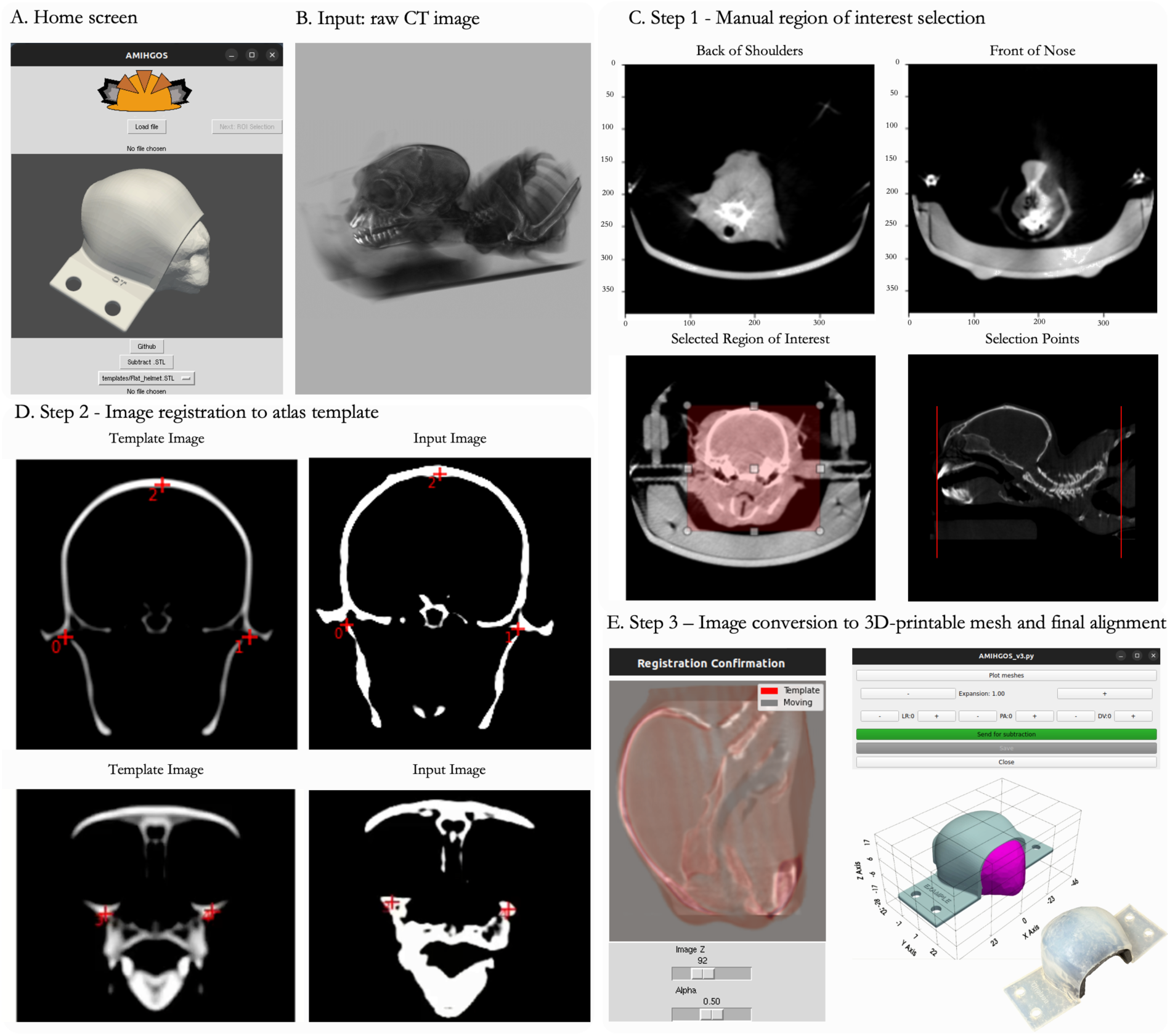
Automated Marmoset Imaging Helmet Generation, an Open-Source Software (AMIHGOS). (A) The software home screen allows users to load NIFTI or DICOM format files as (B) raw CT image inputs. (C) In the ROI selection step, users select boundary points on the nose and just posterior to the glenohumeral joint (just caudal to the shoulder joint) and define the ROI to be included in the helmet generation. (D) Users then align their input image with the template image by selecting corresponding points on the ear canals, the top of the head, and the front of the orbitals in each image. (E) Registration is confirmed by dragging the “Image Z” or “Alpha” bars with the cursor to ensure the template and input images are aligned. (F) Users can adjust any necessary translational modifications and generate an STL file of the customized helmet, which can be sent for 3D printing.

#### 2.3.2 AMIHGOS Step 1: Manual region of interest selection

After loading the image, the user first selects slices at the anterior and posterior bounds of the image (Figure 3C), then draws a region of interest around the head using point-and-click to form the left-right and dorsal-ventral bounds. To accomplish this, we employed the SimpleITK Python library to load and transform CT images^49^. The ROI selection graphical user interface (GUI) was adapted from the SimpleITK Jupyter Notebook tutorial for compatibility with the Tkinter GUI development library^50,51^.

#### 2.3.3 AMIHGOS Step 2: Image registration to atlas template

The image values extracted from Step 1 (binarized Hounsfield Unit values corresponding to the skull/bone) are registered to an averaged CT atlas^3^ image to align the head to a position in which the face is centered in the helmet and the animal can look freely in front of them. We adapted methods from the SimpleITK^49^ to generate an interface for the user to select four or more corresponding fiducials in the atlas (fixed) and uploaded (moving) images (Figure 3D). Through testing, we found that selecting points centered in the interaural canal, the top of the head, and the orbital bone resulted in consistently successful registration to the fixed image. These fiducials are used to calculate the initial affine transformation. The remaining transformations are calculated through optimizing Mattes Mutual Information with gradient descent, both of which are configurations of the SimpleITK ImageRegistrationMethod class^49,50^. The user can confirm accurate registration using an overlaid template and moving images with adjustable position sliders for the Z axis and transparency of the template image. The registered ‘moving file’ of the head in aligned space is then saved.

#### 2.3.4 AMIHGOS Step 3: Image conversion to 3D printable mesh and final alignment

Once the user confirms successful registration (Figure 3E), the image is segmented using methods from dicom2stl^52^. In brief, the registered head is anisotropically smoothed and then thresholded for HU values corresponding to those of skin. The head is then smoothed again with a 15x15x15 voxel median filter, and 5 voxel zero padding is added to the edges of the image. A surface mesh is then extracted and cleaned using VTK’s CleanPolyData filter, small parts are removed, and the number of faces on the head is reduced by 97 % with VTK’s DecimatePro^53^. This processed head mesh is then saved as a STL file. Finally, the user is brought to a mesh manipulation GUI in which a selected template helmet file and head STL file are aligned. By default, the head mesh is expanded along each axis by 15 % to allow room for foam inserts. The user can translate the head along all axes for optimal fit and increase or decrease the Y-axis expansion of the head (Figure 3F). Once the user confirms the mesh alignment, a Boolean difference is calculated between the head and helmet meshes. The resulting helmet then matches the topology of the animal’s head. Sharp edges are smoothed with a Taubin filter, a label is added with an animal identifier, and the final helmet file is saved as a STL file.

#### 2.3.5 Head positioning for MRI signal optimization

Through iterating helmet designs for different head shapes, we have found that contouring the helmet to the brow ridge allows for better signal quality than if the 3D printed helmet and foam are not touching that part of the head. Previously, we corrected such susceptibility-related artifacts^32^ by filling skin-to-air interface gaps with a water-based lubricating gel. Here, we tested the effect of helmet positioning on susceptibility-related artifacts using a marmoset-specific MRI phantom (described and available through the online portal) and helmets conforming (or not) to the brow ridge. An 8-channel receive array was used for these assessments, detailed in the following section.

#### 2.3.6 AMIHGOS testing and validation

Before awake neuroimaging (the CT inputs to AMIHGOS were acquired under 2 % isoflurane anesthesia), marmosets were trained according to established protocols. As detailed in Silva et al. 2011^46^ and later updated in Schaeffer et al. 2021^54^, marmosets were acclimatized to awake MRI in three phases. In phase 1, marmosets were placed in a body restrainer (‘Universal Restrainer’ from the online resource) for increasingly long periods of time, starting at 15 minutes and progressing up to an hour over the course of a week. The marmosets were rewarded at the start and end of every session. In phase 2, marmosets were acclimated to the periodic sounds of the MRI while in a tube similar in size as the scanner inner diameter (120 mm). Performance was tracked using the Behavioral Response Scale^46^, and if a score of 2 or better was achieved (corresponding to mostly quiet, agitated only initially), the marmosets proceeded to phase 3, helmet fitting, and exposure to MRI sounds. During phase 3, scores were continuously monitored (shown in the Results Section below for 46 marmosets) after helmet fitting, and the marmosets proceeded to imaging once reaching a score of 1 or 2. After training, the marmosets proceeded to awake functional MRI or PET experiments, for which the resultant image quality and measured head motion are reported below. We then performed a comparative analysis of helmet motion between a standard foam-lined helmet (1 mm wall thickness without animal-specific contours) and the custom helmets generated by AMIHGOS. To estimate translational and rotational motion, linear registration to the middle volume was conducted using AFNI’s (Analysis of Functional NeuroImages) 3dvolreg^52^.

### 2.4 Conformal radiofrequency coils and supporting hardware for marmoset MRI

Although the marmoset skull is similar in size to that of a rat, their skull and brain morphologies differ significantly^21,55,56^ (Figure 4A). These differences, namely the curvature of the marmoset skull and depth of the temporal lobes, make surface arrays designed for rodents less effective for use with marmosets. Our groups have invested significant effort in developing custom MRI coils for marmosets ^29,39^; currently, the MNSA resource has engineering designs for 12 such coils^29,30,32,39,57^. Here, we describe the design and testing of two recently developed coils: an 8- channel receive array optimized for awake fMRI (to be coupled with AMIHGOS-generated helmets) and a 14-channel coil designed for anatomical imaging in stereotactic position. Both coils were designed in SolidWorks (Version 2022, Dassault Systèmes SolidWorks Corporation, Waltham, MA), along with a thin-walled restrainer (which minimizes multimodal attenuation for PET and CT scans) that is MRI compatible and a 3D printable MRI stereotax device for presurgical alignment (also works for PET/CT scans). We also developed a marmoset body phantom for RF coil assessment and sequence optimization. Using a whole-body CT scan of a 350 g adult marmoset, we created a mesh, imported it into CAD (SolidWorks), and lofted it across body planes to generate a smooth, hollow part that can be filled with phantom solution (H₂O bidistilled water, CuSO₄ 5H₂O 1 g/l, and NaCl 3.6 g/l; Bruker BioSpin MRI GmbH, Ettlingen, Germany). Our restrainer and full-body phantom are suitable for both MRI and CT scans. They were used to characterize our coils and securely hold the marmosets with their AMIHGOS-generated helmets using a localizer RARE (Rapid Acquisition with Relaxation Enhancement) sequence (TR = 100 ms, TE = 8.319 ms, flip angle = 30°, slice thickness = 1 mm, matrix = 256 × 256, FOV = 64 × 64 mm², resolution = 0.250 × 0.250 mm, bandwidth = 10,000 Hz, and a scan time of 12.8 s), as shown in Supplementary Figure 1. Both can be 3D printed with semi-flexible materials such as Durable Resin or Nylon 12 (FormLabs, Somerville, MA, USA), which are MRI-compatible.

**Figure 4:**
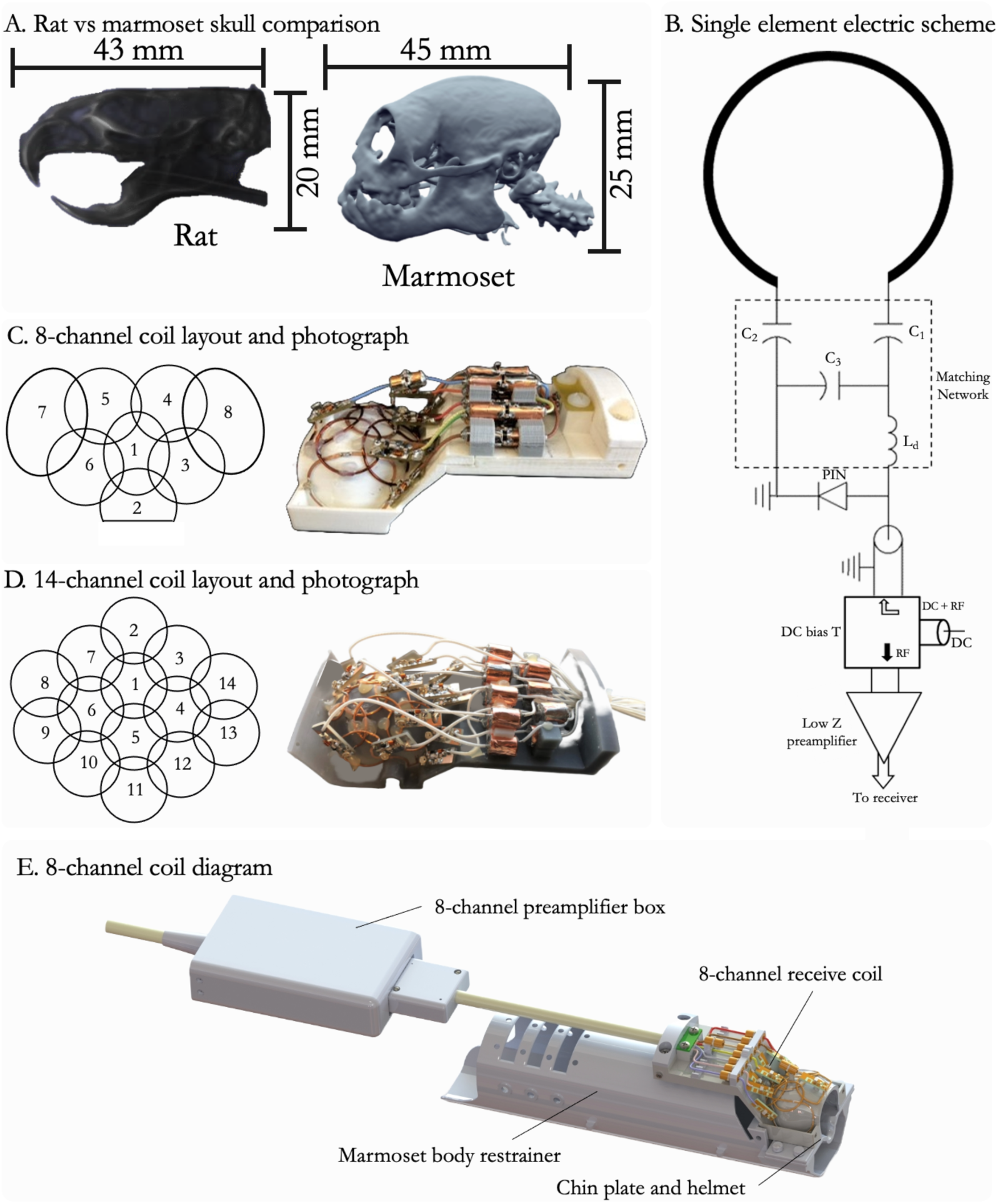
(A) Comparison of the size and morphology of a rat and a marmoset skull. (B) Single-element electrical scheme for a coil, adapted from Papoti et al. 2017^39^. (C) 8-channel coil layout and rendering on a marmoset skull. (D) 14-channel coil layout on a marmoset skull and photograph of assembled coil. (E) Assembly of all components needed for awake imaging with the 8-channel coil, including the coil itself, preamplifier box, body restrainer, chin plate and helmet.

#### 2.4.1 8-channel receive-only radiofrequency array for awake MRI

##### 2.4.1.1 Design and Integration

The radiofrequency receive coil support must be positioned as close as possible to the marmoset’s head to ensure whole brain coverage while maximizing sensitivity during MRI acquisition^29,58^. To fulfill these requirements, the support (‘coil-former’) for the coil elements was designed following the external surface of a standard helmet (external surface of AMIHGOS helmet; Figure 3F) with a wall thickness of 1 mm. Thus, the coil can be placed atop the AMIHGOS helmet with a transition fit, just tight enough to hold it securely in place (Figure 4). While the coil array described here consists of 8 receive channels (Figure 4C), this design allows for flexibility in the number of elements, enabling optimal utilization of the available channels in the user’s MRI scanner. Since marmosets have thick temporal muscles, the receive elements near the temporal lobes were designed with a larger diameter to provide deeper sensitivity into the brain compared to those located over the frontal and parietal lobes.

##### 2.4.1.2 Signal Optimization

Each element of the phased array receive coil was partially overlapped with its nearest neighbors to minimize coupling between elements. This was validated by measuring the S_21_ parameter using a vector network analyzer (ZNL3; Rohde and Schwarz GmbH & Co. KG, Munich, Germany), with the coils tuned to 400 MHz and matched to 50 Ω, while all other elements were detuned. All receive elements were tuned and matched according to the balanced matching network circuit (Figure 4B). During RF transmission, all elements were actively detuned by inserting a resonant inductor (L_d_) with a PIN diode (*MA4P7470F-1072T, MACOM*) biased by +3.6 V/100 mA DC delivered through the same coaxial cable. To prevent coupling with non-nearest neighbor elements, each coil was connected to a low input impedance preamplifier (WMA9RA, WanTcom Inc., Chanhassen, MN) by a λ/2 electrical length cable composed of coaxial cables and phase shifters.

Due to space constraints near the coil elements, the preamplifiers were housed in a box located 200 mm away from the coil end. These preamplifiers were mounted on two printed circuit boards, each accommodating four preamplifiers. Each coil element was connected to the preamplifiers using 50 Ω RG-196 coaxial cables (RG-196; Leoni Wiring Systems, Inc., Lake Orion, MI, USA) equipped with 50 Ω (ODU MAC) coaxial connectors. These connectors were assembled into a custom 2 x 4 connector case, designed and 3D printed (Pro3; Raise3D Technologies, Irvine, CA, USA) in-house with polylactic acid (PLA).

##### 2.4.1.3 Testing and Validation

The performance of the receive elements was first characterized at the workbench by measuring the quality factor when loaded and unloaded with the marmoset phantom. To assess the isolation provided by the active detuning circuit and the low impedance preamplifiers, the S_21_ parameter was measured when the coil was terminated to 50 Ω and compared to when the elements were connected to the preamplifiers. Individual signal-to-noise ratio (SNR) maps and noise correlation matrices were acquired in the 9.4 T scanner with the coil loaded with the marmoset phantom described above. In vivo, MRI from awake marmosets was acquired using all the hardware components described here. For these experiments, a 120 mm inner diameter 16-rung quadrature-driven Birdcage coil (Resonance Research, Billerica, MA) with active detuning was used to characterize the 8-channel receive coil.

#### 2.4.2 14-channel receive-only radiofrequency array for anesthetized MRI

##### 2.4.2.1 Design and Integration

A 14-channel receive-only array was developed and optimized for a 9.4T/30 cm ultra- shield refrigerated magnet (Bruker Biospin Inc, Ettlingen, Germany). Each coil element was constructed using AWG-18 copper wire, with most elements having a 20 mm diameter. Notably, the elements adjacent to the temporal lobe (coils #8, 9, 13, and 14) were designed with a 25 mm diameter to enhance penetration and brain sensitivity, accommodating the marmoset’s thick temporalis muscle. The support structure for assembling the coil was designed in SolidWorks (Version 2022, Dassault Systèmes SolidWorks Corporation, Waltham, MA) and 3D printed in gray resin (Formlabs, Somerville, MA). The array (Figure 4D) is designed to accommodate the placement of ear bars for stereotactic alignment when the animal is anesthetized and allows for a face mask to deliver inhalant anesthesia (e.g., isoflurane).

##### 2.4.2.2 Signal Optimization

To minimize mutual inductance, all nearest neighbor elements were partially overlapped. The coil circuitry for each element in the 14-channel receive array consisted of a matching network and a PIN diode-controlled blocking circuit for active detuning^59^. To avoid common modes in the cables, cable traps consisting of 3-turns with the receive coaxial cables were inserted between the matching network and the preamplifiers. Decoupling between non-nearest neighbor coils was achieved by connecting the elements to low-noise preamplifiers through a pi-network phase shifter, combined with 50 Ω coaxial cables (RG-196) adjusted to provide an equivalent λ/2 cable at the input of the preamplifiers (WMA9RA, WanTcom Inc., Chanhassen, MN). The preamplifiers were assembled in four printed circuit boards (PCBs), each supporting four channels. All the loop elements, matching network, and active detuning circuits were integrated into the 3D printed support structure.

##### 2.4.2.3 Testing and Validation

The coil’s performance was characterized on the workbench^40^ using a vector network analyzer. This evaluation included the unloaded/loaded quality factor (Q) ratio assessment and measured the isolation provided by the active detuning circuit and the preamplifier decoupling. Individual SNR maps and noise correlation matrices were obtained from the MRI scanner with the coil loaded with a phantom filled with CuSO_4_ x 2H_2_O 1g/L. T2 weighted images with 250 µm isotropic resolution were acquired in a 9.4T MRI scanner – as described in the Methods Section – connected to an Avance Neo console running ParaVision 360. The imaging sequence used was a T2-weighted RARE with TE = 48 ms, TR = 7209 ms, RARE factor = 7, matrix = 256 x 256, FOV = 40 x 40 mm, slice thickness = 0.5 mm and five averages, resulting in acquisition time of 21 min 37 s.

## 3. Results

### 3.1 AMIHGOS Validation

Using AMIHGOS, custom-fitted helmets were generated from CT images for awake PET ([¹⁸F] Fluorodeoxyglucose at 80 MBq, 60 min static acquisition) and MRI scans. This process ensured that each marmoset received a helmet tailored to its specific anatomy, thereby optimizing the fit and stability during imaging. Figure 5A illustrates the progression through awake restraint training for 46 marmosets, with helmet fitting taking place on day 10. Each day, we recorded their comfort levels using a Behavioral Response Scale^46^, where a score of 8 indicated agitation and stress, and a score of 1 indicated calmness and relaxation^46^. Initially, the marmosets exhibited an activity level of approximately 4, but this quickly decreased as they became more accustomed to the MRI-specific environmental conditions (e.g., body restrainer, MRI sounds, and helmet) – by day 14, the average score had reduced to 1 (Figure 5A). Figure 5B shows successful imaging outcomes across different modalities (PET, MRI T2, and MRI EPI) for a ‘small’ (325 g), ‘medium’ (425 g), and ‘large’ (500 g) adult marmosets using AMIHGOS generated helmets. We also compared helmet motion between a universal foam-lined helmet (with a 1 mm wall thickness but no animal-specific contours) and the AMIHGOS-generated custom helmets. Figure 5C demonstrates that the custom helmets significantly reduced translational and rotational motion, as estimated by linear registration to the middle volume. This reduction in translational and rotational head movement was achieved without requiring invasive methods such as surgically implanted chambers, ensuring that the marmosets remained comfortable and secure throughout the scanning process. The AMIHGOS helmets thus provide a reliable, non-invasive solution for minimizing motion artifacts in awake imaging sessions.

**Figure 5:**
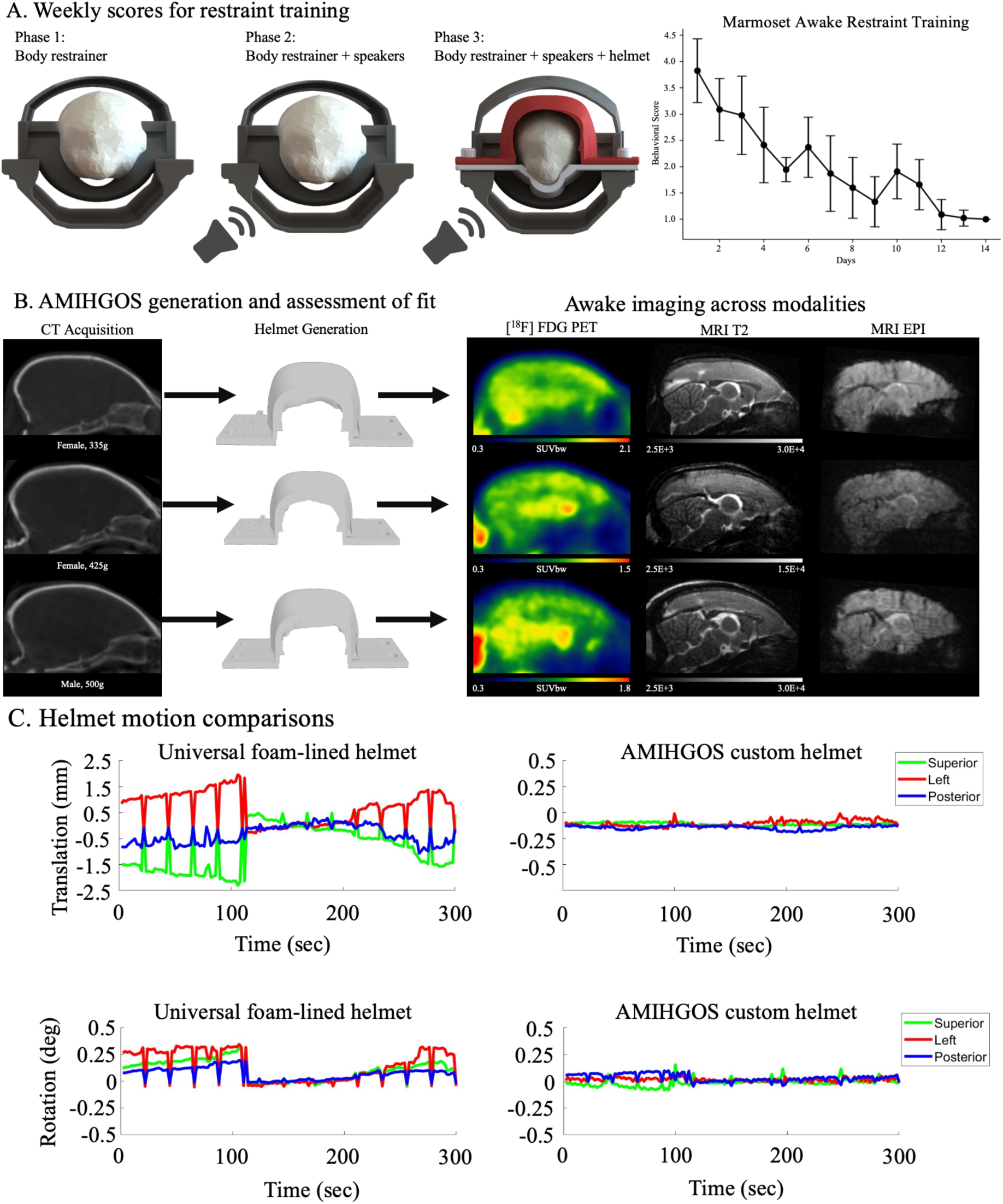
(A) Restraint training progression over 14 days, showing the phases of adaptation to the MRI environment. (B) CT images were used to generate custom-fitted helmets tailored to each marmoset’s anatomy, ensuring an optimized fit for stable, awake PET ([^18^F] FDG, 80 MBq, 60-minute static acquisition), PET, and MRI scans, showing fit for small (325 g), medium (425 g), and large (500 g) adult marmosets. (C) Helmet motion comparison between a universal foam-lined helmet (1 mm wall thickness) and AMIHGOS-generated custom helmets.

### 3.2 Conformal radiofrequency coils for MRI

#### 3.2.1 8-channel receive-only radiofrequency array

The 8-channel receive-only coil was characterized on the workbench and tested on a 9.4T 30 cm horizontal bore MRI scanner (Bruker BioSpin Corp, Billerica, MA) with the previously described marmoset whole body phantom. Coil performance was assessed by acquiring noise-only images (i.e., with no radiofrequency power) and calculating the noise correlation matrix. To evaluate the coil performance during GeneRalized Autocalibrating Partial Parallel Acquisition (GRAPPA) accelerated images, (geometry) g-factor maps were obtained with both anterior-posterior and left-right in the phase encoding directions.

#### 3.2.1.1 Workbench characterization

The 8-channel coil underwent extensive testing and characterization on the workbench using a vector network analyzer with a pair of decoupled pick-up loops built from semirigid coaxial cables. The quality factor of a typical unloaded loop element was Q_u_ = 223, while the loaded quality factor was Q_L_ = 134, resulting in a ratio Q_u_/Q_L_ = 1.6. Such a ratio indicates that the coil elements operate in a coil noise-dominated regime with efficiency and sensitivity in minimizing sample noise, consistent with findings from prior studies^39,60^. The average S_11_ measurement for each channel, with all other channels actively detuned, resulted in 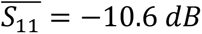. Channel isolation, achieved through partial overlap between nearest neighbor elements, averaged -15.8 dB. Decoupling through the low input impedance preamplifier provided isolation greater than 15 dB across all channels. The active detuning circuit, controlled by PIN diodes, achieved an average, 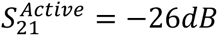.

##### 3.2.1.2 Correlation Matrix, SNR maps, and g-factor maps

The noise correlation matrix (Figure 6A) shows that most channels exhibit low noise correlations, with a mean correlation coefficient of 17.4%. The minimum correlation is 2.0%, and the maximum reaches 47.0%, likely due to residual coupling between channels 4 and 8. The standard deviation of 11.8% reflects moderate variability across the correlations, indicating some uneven distribution, which could be attributed to the larger coil placed around the temporalis muscles – leading to increased sensitivity. The low average noise correlation suggests that the channels are largely independent (uncoupled), which increases the image quality and allows for parallel imaging.

**Figure 6:**
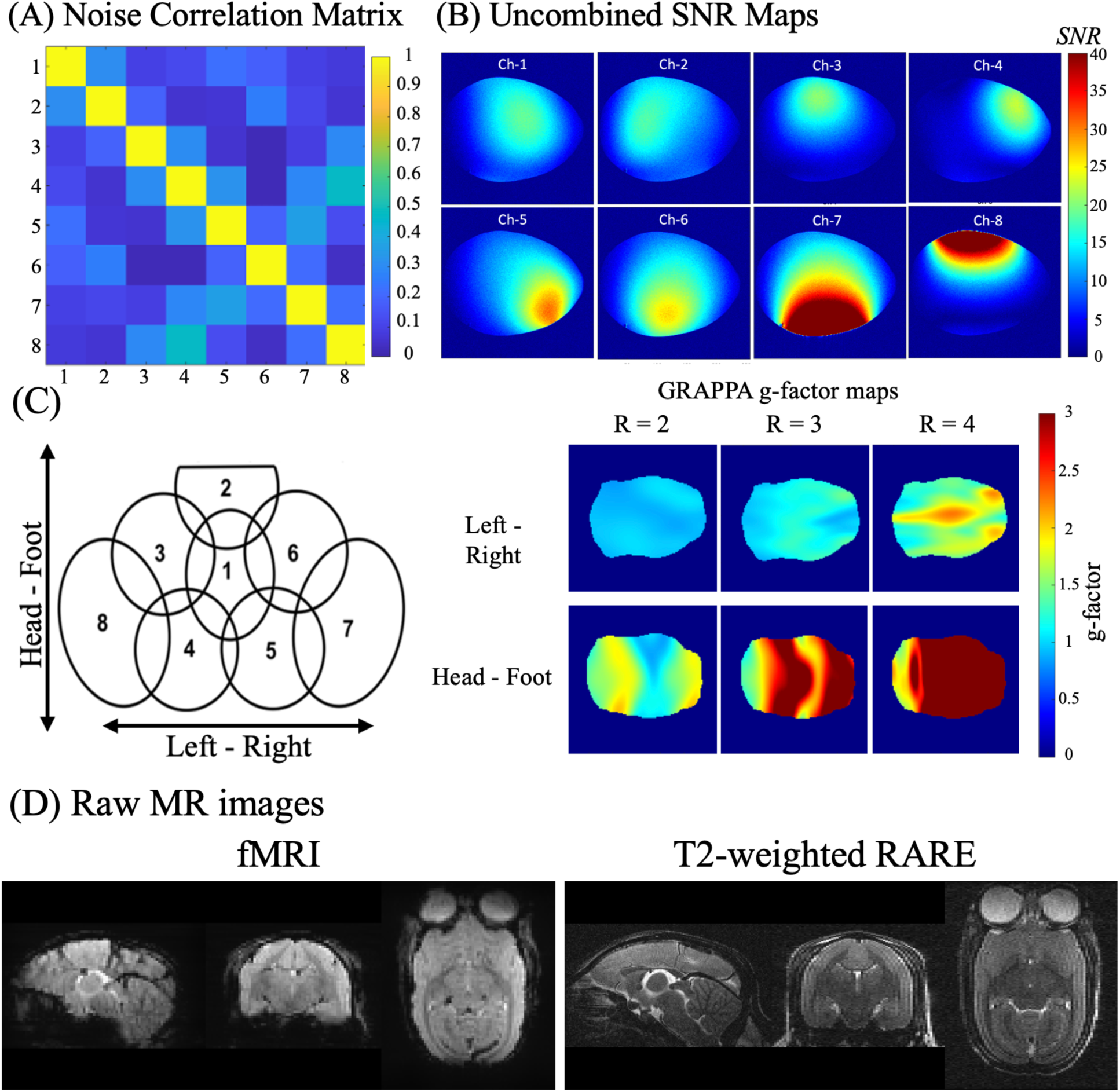
(A) Noise correlation matrix showing the correlation levels between channels, with an average correlation coefficient of 10.7% across channels. (B) Uncombined SNR maps from the central coronal plane demonstrate individual coil element sensitivity. (C) Schematic of the 8-channel coil layout and GRAPPA g-factor maps. (D) Raw MR images from a marmoset brain, including fMRI and T2-weighted RARE sequences.

Figure 6B displays the uncombined SNR maps acquired from the central plane in the coronal orientation, highlighting the high sensitivity of each element according to the coil layout shown in Figure 6C. Notably, elements 7 and 8 exhibit deeper sensitivity penetration, particularly in the temporal lobes where the marmoset’s temporalis muscles create a thick layer. The uncombined SNR maps indicate effective decoupling between each element and its nearest neighbors, achieved through partial overlap and decoupling provided by the low input impedance preamplifiers.

The noise amplification due to GRAPPA accelerated acquisition is quantified by the g-factor maps shown in Figure 6C. These maps suggest that the optimal direction for acceleration is left-right, with a feasible acceleration factor of 2.

#### 3.2.2 14-channel receive-only radiofrequency array

##### 3.2.2.1 Workbench characterization

Using the methods described above, network analyzer measurements showed that the isolation provided by the PIN diode-controlled active detuning circuit averaged 32 dB. The low input impedance preamplifiers offered isolation better than 15 dB across all elements. The unloaded quality factor (Q_u_) was measured at 253, and the loaded quality factor (Q_L_) was 150. This resulted in a Q_u_/Q_L_ ratio of 1.7, indicating that the coil operates primarily in a coil noise-dominated regime. Figure 7A presents the noise correlation matrix derived from a noise-only acquisition. The average correlation coefficient for the non-diagonal elements was 8.3%, with a maximum correlation of 27.2% observed between channels 6 and 10, indicating residual coupling between these channels due to the higher number of elements.

**Figure 7:**
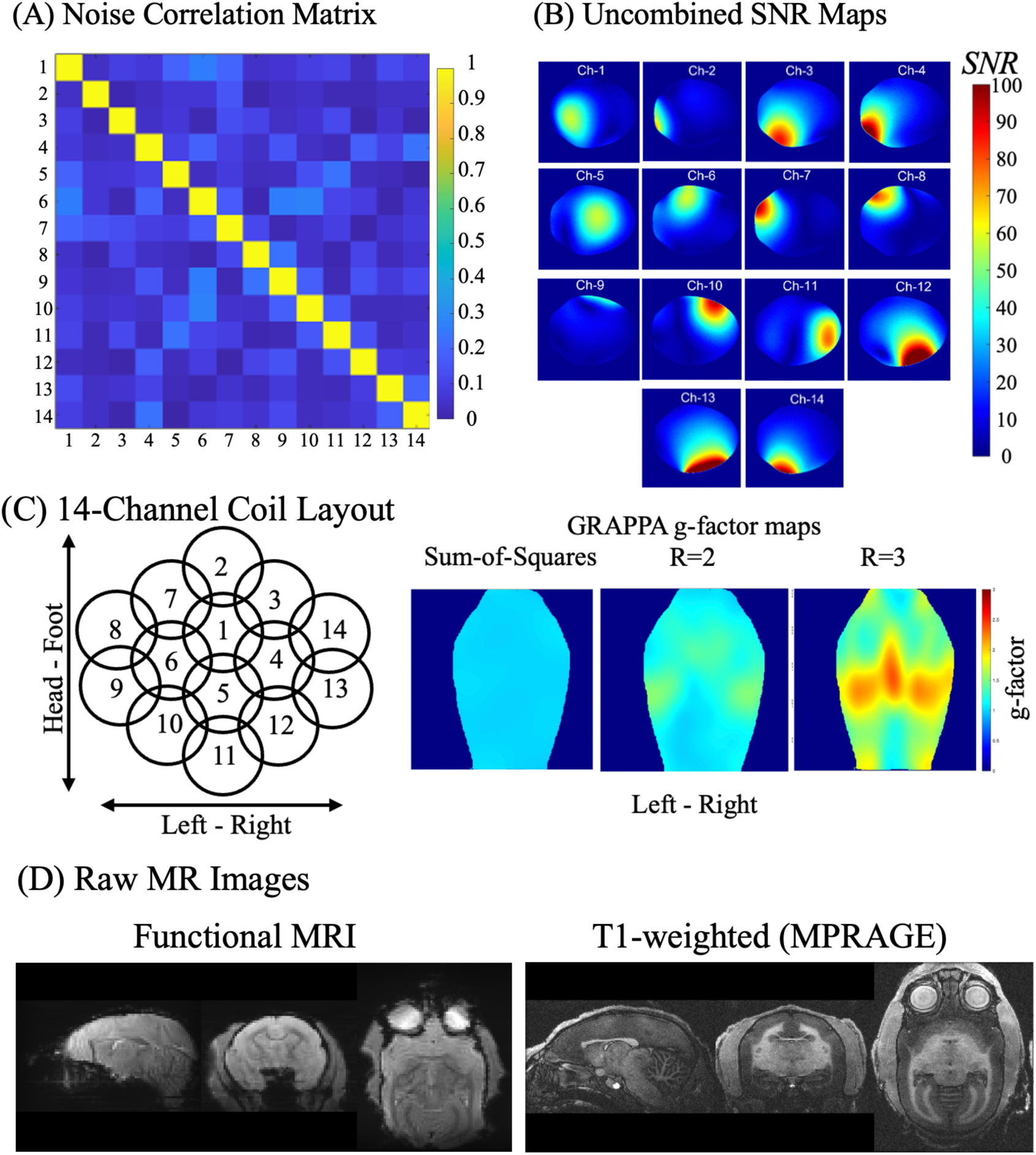
(A) Noise correlation matrix from a noise-only acquisition, showing low inter-channel correlations with an average of 8.3% and a maximum correlation of 27.2% between channels 13 and 14, indicating minimal residual coupling. (B) Uncombined SNR maps from a central coronal orientation, demonstrating individual coil element sensitivity. (C) Layout of the 14-channel coil array and GRAPPA g-factor maps. (D) Raw fMRI and T1-weighted MPRAGE MRI acquired from a marmoset.

##### 3.2.2.2 Correlation Matrix, SNR maps, and g-factor maps

The individual SNR maps obtained from a coronal orientation of a phantom are illustrated in Figure 7B. While most elements demonstrate effective decoupling, channel 9 shows a noticeable loss of sensitivity. Despite this, the sum-of-squares reconstructed image shows robust coverage with high sensitivity throughout the entire volume of interest. The T1-weighted anatomical brain images of a marmoset, presented in Figure 7D, were acquired with an in-plane resolution of 156 µm and a slice thickness of 500 µm. The noise correlation matrix for the 14-channel coil yielded a mean value of 8.3%, with a standard deviation of 6.7%, a maximum correlation of 27.2%, and a minimum correlation of 0.02%.

The Q measurements for the loaded and unloaded condition indicate that, for this element size and frequency of operation, the coil losses are mainly in the coil dominating regime. The SNR maps indicate that most coil elements are well decoupled and maintain high local sensitivity, as confirmed by the noise correlation matrix.

## 4. Discussion

The goal of the Marmoset Neuroscientific Apparatus (MNSA) resource is to centralize the engineering efforts that have supported over a decade of marmoset neuroscience in a publicly available repository. In parallel with our efforts to make our neuroimaging data publicly available through the Marmoset Brain Connectome (MBC) and following in the footsteps of open-science initiatives^61^, we hope to set a new standard of sharing the methodological details that are often obfuscated in published works and remain as knowledge within individual laboratories. The scientific value of the marmoset model has been well established, resulting in increased global adoption; however, custom apparatus remains necessary for many experimental applications. With insufficient tools and resources tailored to marmoset research, our team has compiled thousands of hours into designing, developing, and rigorously testing a wide array of tools, hardware, software, and apparatus specific to marmoset neuroscience to address this gap. These resources (exemplars shown in Figures 1 and 2) are now freely accessible through our online platform (https://www.marmosetbrainconnectome.org/apparatus/), designed to accelerate advancements in the field by providing accessible and reproducible resources. This manuscript discusses the design, development, and validation of three such resources.

With functional imaging methods such as functional MRI or PET offering the distinct advantage of being entirely noninvasive while also showing whole-brain dynamics, a major goal of our work over the past decade has been to develop the means to conduct these modalities in fully awake, behaving marmosets. Early designs included a conformal helmet and a jacket worn by the marmoset that was then mounted to a suspended chassis system^38,62^. Designs to be used in concert with head chambers for electrophysiology were also developed, allowing for near- motionless functional MRI^32,35^. As training strategies evolved for studies that required longitudinal imaging of large cohorts (200+) of marmosets, we sought to optimize and automate the development of conformal head-fixation devices. For this reason, we generated AMIHGOS, a user-friendly software that uses a CT image of a marmoset head to automatically generate individualized helmets based on each animal’s unique morphology. The latest version of the software is made available through the online portal. In Figure 5, we demonstrate the training outcomes of these conformal helmets when used with our optimized body-restraint system (optimized for body size across the age span, minimal image attenuation, and minimal physical interaction with the marmoset to avoid undue stress; Figure 5A). Figure 5B demonstrates the resultant imaging outcomes across PET, structural, and functional MRI after the animals were fitted with a helmet and well-trained. It is critical to note that training is just as crucial as a well- conforming helmet – a calm and comfortable marmoset is less likely to move during imaging sessions. Figure 5C shows an example of < 100 µm of translational head motion during awake functional MRI from a well-fitted AMIHGOS-generated helmet.

The conformal helmets were designed for multimodal imaging in the sphinx position (e.g., Figure 1B) because this body position is amenable to a variety of awake functional task-based designs. For example, this position allows them to see a screen for visual experiments, lick a reward tube, or be presented with an olfactory stimulus. Here, we present an 8-channel radiofrequency array that slips on over any marmoset helmet and allows for functional MRI at 9.4T. The CAD designs for this coil and, at present, a dozen others are available through the online resource. Figure 4 shows the coil layout and circuitry support of this coil system. In contrast, Figure 6 demonstrates benchtop validation of decoupled elements with high SNR (Figure 6A & B), capable of accelerated functional imaging (Figure 6C). Figure 6D illustrates the mean image from the raw functional MRI data acquired fully awake over 20 minutes. Similarly, this coil can be used to acquire high-fidelity anatomical images, with Figure 6D showing a T2w RARE image acquired from an awake marmoset. Both such images are free of any significant motion artifacts. Unavoidable movements such as breathing (and sniffing), eye movements, and heartbeat can be further ameliorated with image regression if those signals can be reliably recorded. Supplementary Figure 1 demonstrates some of the iterative tuning that we have conducted to optimize the position of the head relative to the helmet system. With the goal of not impeding vision or compressing the skin around the eyes, we needed the helmet to be a few millimeters posterior to the anterior extent of the brow ridge. We found, however, that if the former did not tightly contour the dorsal surface of the brow, susceptibility-related artifacts were present in the fMRI images (due to a transition from skin to air). As shown in Supplementary Figure 1, we recovered this signal through optimized head placement in AMIHGOS and tight contouring of foam within the helmet.

In parallel with the awake imaging coil development, we have developed a variety of coils for anesthetized imaging in marmosets^30,63^, mainly for longer sequence schemes that do not require the animal to be awake (e.g., T1, T2, diffusion contrasts). One benefit of these coil schemes is that elements can be placed more liberally across the head morphology (see Figure 4D), unencumbered by the need for a clear path for vision or the chin-plate of our helmet system. When used in concert with a stereotactic position system compatible with PET and CT imaging, rigid mechanical alignment can be readily transferred to a surgical stereotaxis when presurgical anatomical or functional alignment is necessary. Such mechanical designs are freely available through the online resource. Figure 7 demonstrates well-decoupled elements in a 14-channel scheme with a high signal-to-noise ratio, as shown in the raw images in Figure 7D. Depending on the capabilities of individual MRI systems (namely the number of receiving channels), the channel count can be altered and optimized to fit atop this ‘coil-former’, with several designs available through the online resource.

Although it is not possible to discuss all designs available on https://www.marmosetbrainconnectome.org/apparatus/ within the scope of a single manuscript, we also make available designs for other modalities, including tools for histology, a variety of behavioral apparatus (including complete touchscreen systems, stimuli, and training standard operating procedures), a collection of 3D printable head chambers for electrophysiology or optical imaging, among many other designs. Our functional MRI task designs are also available, down to the stimuli, with preprocessing code made available through marmosetbrainconnectome.org. With increased attention to social experiments in marmosets^13,45,64^, we also make engineering designs for multi-marmoset fMRI and PET available through the portal.

These many designs made available here are by no means comprehensive, with many other groups across the globe having developed marmoset neuroscientific apparatuses that, in one way or another, have inspired what we present here. Our goal here was to provide a publicly available platform through which the community can house their designs. With explicit instructions for contributions at https://www.marmosetbrainconnectome.org/apparatus/submissions.html, we hope to grow the resource to be representative of the full breadth of the burgeoning field of marmoset neuroscience. Indeed, the availability and support of public resources have been nothing less than impressive over the past decade^1–12^. One area in particular, optical imaging, offers great promise in marmosets compared to Old World primates^65–69^. The lissencephalic cortex of the marmoset allows for imaging areas of the cortex (e.g., dorsolateral prefrontal area), unencumbered by sulci as in Old World primates like macaques. Among many others, the category of optical imaging is currently underrepresented in the resource; thus, such designs are welcomed from the field. An important detail regarding attribution is that if the designs were used to collect data for a previously published paper but were not explicitly represented in that manuscript (to the extent permitted by the publisher), a citation will be included to attribute the resource, as illustrated in Figure 1B.

## Acknowledgments

We thank Lauren Dubberley, Brianne Stein, Kevin Thiel, and Dr. Julia Oluoch at the University of Pittsburgh for animal care and preparation. This work was supported by the National Institute of Neurological Disorders and Stroke of the National Institutes of Health under Award Number R21NS125372, as well as the National Institute on Aging of the National Institutes of Health, under Award Numbers U19AG074866 and R24AG073190. We also thank the Hearing and Vision Sciences Graduate Certificate Program. Work performed at the University of Western Ontario was supported by the Canada Foundation for Innovation, Canada First Research Excellence Fund to BrainsCAN, Brain Canada Platform Support Grant, Canadian Institutes of Health Research, and a Discovery grant by the Natural Sciences and Engineering Research Council of Canada.

---

**Supplementary Figure 1.**
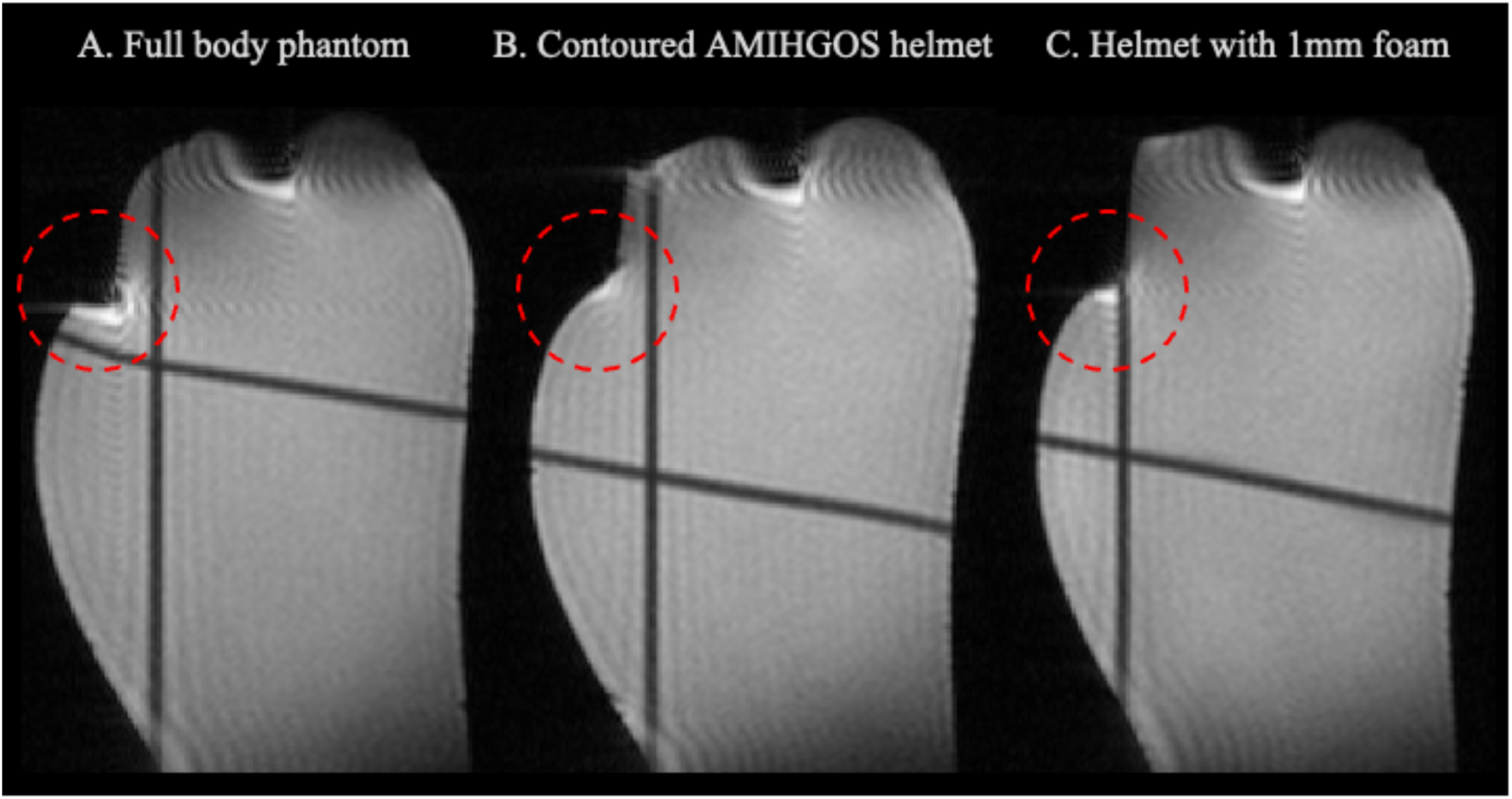
(A) Phantom without a helmet, used to visualize existing susceptibility artifacts. (B) 3D printed durable helmet covering the sagittal bump of the phantom corresponding to the brow ridge (red circle). (C) Helmet with an added 1mm foam layer on the inner side, further minimizing susceptibility artifacts. The susceptibility artifact is reduced with the addition of the AMIHGOS-generated helmet, as the interface between the phantom and air is replaced by foam and plastic materials with magnetic susceptibilities closer to that of biological tissue.

## References

1. Gilbert, K. M. et al. Open-source hardware designs for MRI of mice, rats, and marmosets: Integrated animal holders and radiofrequency coils. J Neurosci Methods 312, 65–72 (2019).

2. Kita, Y. et al. Cellular-resolution gene expression profiling in the neonatal marmoset brain reveals dynamic species- and region-specific differences. Proceedings of the National Academy of Sciences 118, (2021).

3. Liu, C. et al. Marmoset Brain Mapping V3: Population multi-modal standard volumetric and surface-based templates. Neuroimage 226, 117620 (2021).

4. Liu, C. et al. A resource for the detailed 3D mapping of white matter pathways in the marmoset brain. Nat Neurosci 23, 271–280 (2020).

5. Liu, C. et al. A digital 3D atlas of the marmoset brain based on multi-modal MRI. Neuroimage 169, 106–116 (2018).

6. Majka, P. et al. Open access resource for cellular-resolution analyses of corticocortical connectivity in the marmoset monkey. Nat Commun 11, 1133 (2020).

7. Majka, P. et al. Towards a comprehensive atlas of cortical connections in a primate brain: Mapping tracer injection studies of the common marmoset into a reference digital template. Journal of Comparative Neurology 524, 2161–2181 (2016).

8. Okano, H. et al. Brain/MINDS: A Japanese National Brain Project for Marmoset Neuroscience. Neuron 92, 582–590 (2016).

9. Schaeffer, D. J. et al. An open access resource for functional brain connectivity from fully awake marmosets. Neuroimage 252, 119030 (2022).

10. Shimogori, T. et al. Digital gene atlas of neonate common marmoset brain. Neurosci Res 128, 1–13 (2018).

11. Tian, X. et al. An integrated resource for functional and structural connectivity of the marmoset brain. Nat Commun 13, 7416 (2022).

12. Zhu, X. et al. An anatomical and connectivity atlas of the marmoset cerebellum. Cell Rep 42, 112480 (2023).

13. Miller, C. T. et al. Marmosets: A Neuroscientific Model of Human Social Behavior. Neuron 90, 219–233 (2016).

14. Okano, H., Hikishima, K., Iriki, A. & Sasaki, E. The common marmoset as a novel animal model system for biomedical and neuroscience research applications. Semin Fetal Neonatal Med 17, 336–340 (2012).

15. Sukoff Rizzo, S. J., et al. Bridging the rodent to human translational gap: Marmosets as model systems for the study of Alzheimer’s disease. Alzheimer’s & Dementia: Translational Research & Clinical Interventions 9, (2023).

16. Homanics, G. E. et al. Early molecular events of autosomal-dominant Alzheimer’s disease in marmosets with PSEN1 mutations. Alzheimer’s & Dementia 20, 3455–3471 (2024).

17. Ando, K. et al. PET Analysis of Dopaminergic Neurodegeneration in Relation to Immobility in the MPTP-Treated Common Marmoset, a Model for Parkinson’s Disease. PLoS One 7, e46371 (2012).

18. Roe, A. W., Fritsches, K. & Pettigrew, J. D. Optical imaging of functional organization of V1 and V2 in marmoset visual cortex. Anat Rec A Discov Mol Cell Evol Biol **287A**, 1213– 1225 (2005).

19. Schaeffer, D. J. et al. Diffusion-weighted tractography in the common marmoset monkey at 9.4T. J Neurophysiol 118, 1344–1354 (2017).

20. Schaeffer, D. J. et al. Face selective patches in marmoset frontal cortex. Nat Commun 11, 4856 (2020).

21. Ogihara, N. et al. Muscle architectural properties in the common marmoset (Callithrix jacchus). Primates 58, 461–472 (2017).

22. Matsuzaki, M. & Ebina, T. Common marmoset as a model primate for study of the motor control system. Curr Opin Neurobiol 64, 103–110 (2020).

23. Mitchell, J. F. & Leopold, D. A. The marmoset monkey as a model for visual neuroscience. Neurosci Res 93, 20–46 (2015).

24. Elston, G. N., Tweedale, R. & Rosa, M. G. P. Cellular heterogeneity in cerebral cortex: A study of the morphology of pyramidal neurones in visual areas of the marmoset monkey. J Comp Neurol 415, 33–51 (1999).

25. Hung, C.-C. et al. Functional Mapping of Face-Selective Regions in the Extrastriate Visual Cortex of the Marmoset. The Journal of Neuroscience 35, 1160–1172 (2015).

26. Luongo, F. J. et al. Mice and primates use distinct strategies for visual segmentation. Elife 12, (2023).

27. Schaeffer, D. J. et al. Divergence of rodent and primate medial frontal cortex functional connectivity. Proceedings of the National Academy of Sciences 117, 21681–21689 (2020).

28. Prescott, M. J. & Poirier, C. The role of MRI in applying the 3Rs to non-human primate neuroscience. Neuroimage 225, 117521 (2021).

29. Gilbert, K. M. et al. A radiofrequency coil to facilitate task-based fMRI of awake marmosets. J Neurosci Methods 383, 109737 (2023).

30. Gilbert, K. M. et al. A geometrically adjustable receive array for imaging marmoset cohorts. Neuroimage 156, 78–86 (2017).

31. Papoti, D., Yen, C. C., Mackel, J. B., Merkle, H. & Silva, A. C. An embedded four-channel receive-only RF coil array for fMRI experiments of the somatosensory pathway in conscious awake marmosets. NMR Biomed 26, 1395–1402 (2013).

32. Schaeffer, D. J. et al. Integrated radiofrequency array and animal holder design for minimizing head motion during awake marmoset functional magnetic resonance imaging. Neuroimage 193, 126–138 (2019).

33. Hori, Y. et al. Interspecies activation correlations reveal functional correspondences between marmoset and human brain areas. Proceedings of the National Academy of Sciences 118, (2021).

34. Hori, Y. et al. Cortico-Subcortical Functional Connectivity Profiles of Resting-State Networks in Marmosets and Humans. The Journal of Neuroscience 40, 9236–9249 (2020).

35. Schaeffer, D. J. et al. Task-based fMRI of a free-viewing visuo-saccadic network in the marmoset monkey. Neuroimage 202, 116147 (2019).

36. Hori, Y. et al. Altered Resting-State Functional Connectivity Between Awake and Isoflurane Anesthetized Marmosets. Cerebral Cortex 30, 5943–5959 (2020).

37. Liu, J. V. et al. fMRI in the awake marmoset: Somatosensory-evoked responses, functional connectivity, and comparison with propofol anesthesia. Neuroimage 78, 186–195 (2013).

38. Papoti, D., Yen, C. C., Mackel, J. B., Merkle, H. & Silva, A. C. An embedded four-channel receive-only RF coil array for fMRI experiments of the somatosensory pathway in conscious awake marmosets. NMR Biomed 26, 1395–1402 (2013).

39. Papoti, D. et al. Design and implementation of embedded 8-channel receive-only arrays for whole-brain MRI and fMRI of conscious awake marmosets. Magn Reson Med 78, 387–398 (2017).

40. Papoti, D. et al. Optimization of an 8-channel receive-only surface array for whole brain MRI of marmosets. in Proceedings of the 23rd Annual Meeting ISMRM 3176 (Proceedings of the 23rd Annual Meeting ISMRM, 2015).

41. Murai, T. et al. Improving preclinical to clinical translation of cognitive function for aging-related disorders: the utility of comprehensive touchscreen testing batteries in common marmosets. Cogn Affect Behav Neurosci 24, 325–348 (2024).

42. Cope, Z. A., Murai, T. & Sukoff Rizzo, S. J. Emerging Electroencephalographic Biomarkers to Improve Preclinical to Clinical Translation in Alzheimer’s Disease. Front Aging Neurosci 14, (2022).

43. Murai, T. & Sukoff Rizzo, S. J. The Importance of Complementary Collaboration of Researchers, Veterinarians, and Husbandry Staff in the Successful Training of Marmoset Behavioral Assays. ILAR J 61, 230–247 (2020).

44. Dureux, A., Zanini, A. & Everling, S. Face-Selective Patches in Marmosets Are Involved in Dynamic and Static Facial Expression Processing. The Journal of Neuroscience 43, 3477–3494 (2023).

45. Gilbert, K. M. et al. Simultaneous functional MRI of two awake marmosets. Nat Commun 12, 6608 (2021).

46. Silva, A. C. et al. Longitudinal Functional Magnetic Resonance Imaging in Animal Models. in 281–302 (2011). doi:10.1007/978-1-61737-992-5_14.

47. Miocinovic, S. et al. Stereotactic neurosurgical planning, recording, and visualization for deep brain stimulation in non-human primates. J Neurosci Methods 162, 32–41 (2007).

48. Liang, L. et al. An open-source MRI compatible frame for multimodal presurgical mapping in macaque and capuchin monkeys. J Neurosci Methods 407, 110133 (2024).

49. Lowekamp, B. C., Chen, D. T., Ibáñez, L. & Blezek, D. The Design of SimpleITK. Front Neuroinform 7, (2013).

50. Yaniv, Z., Lowekamp, B. C., Johnson, H. J. & Beare, R. SimpleITK Image-Analysis Notebooks: a Collaborative Environment for Education and Reproducible Research. J Digit Imaging 31, 290–303 (2018).

51. Python Software Foundation. Tkinter – Python interface to Tcl/Tk. Python 3 Documentation.

52. Cox, R. W. AFNI: Software for Analysis and Visualization of Functional Magnetic Resonance Neuroimages. Computers and Biomedical Research 29, 162–173 (1996).

53. Schroeder, W., Martin, K. & Lorensen, B. The Visualization Toolkit. (Kitware, Clifton Park, NY, 2006).

54. Schaeffer, D. J., Liu, C., Silva, A. C. & Everling, S. Magnetic Resonance Imaging of Marmoset Monkeys. ILAR J 61, 274–285 (2020).

55. Casteleyn, C. et al. Anatomical description and morphometry of the skeleton of the common marmoset ( Callithrix jacchus ). Lab Anim 46, 152–163 (2012).

56. Greene, E. C. Anatomy of the Rat. Transactions of the American Philosophical Society 27, ii (1935).

57. Papoti, D. et al. Segmented solenoid RF coils for MRI of ex vivo brain samples at ultra-high field preclinical and clinical scanners. J Magn Reson Open 16–17, 100103 (2023).

58. Silva, A. C. Anatomical and functional neuroimaging in awake, behaving marmosets. Dev Neurobiol 77, 373–389 (2017).

59. Griswold, M. A. Characterization of Multichannel Coil Arrays on the Benchtop. in *Encyclopedia of Magnetic Resonance* (John Wiley & Sons, Ltd, Chichester, UK, 2012). doi:10.1002/9780470034590.emrstm1141.

60. Gilbert, K. M. et al. Concentric radiofrequency arrays to increase the statistical power of resting-state maps in monkeys. Neuroimage 178, 287–294 (2018).

61. Milham, M. et al. Toward next-generation primate neuroscience: A collaboration-based strategic plan for integrative neuroimaging. Neuron 110, 16–20 (2022).

62. Papoti, D., Yen, C., Mackel, J., Merkle, H. & Silva, A. C. A whole-brain 8 channel receive-only embedded array for MRI and fMRI of conscious awake marmosets at 7 T. in Proc 21st Annual Meeting and Exhibit of the International Society for Magnetic Resonance and Medicine (ISMRM) 2778 (Salt Lake City, 2013).

63. Gilbert, K. M., Gati, J. S., Barker, K., Everling, S. & Menon, R. S. Optimized parallel transmit and receive radiofrequency coil for ultrahigh-field MRI of monkeys. Neuroimage 125, 153–161 (2016).

64. Cerrito, P., Gascon, E., Roberts, A. C., Sawiak, S. J. & Burkart, J. M. Neurodevelopmental timing and socio-cognitive development in a prosocial cooperatively breeding primate ( *Callithrix jacchus* ). Sci Adv 10, (2024).

65. Song, X. et al. Mesoscopic landscape of cortical functions revealed by through-skull wide- field optical imaging in marmoset monkeys. Nat Commun 13, 2238 (2022).

66. Iwano, S. et al. Single-cell bioluminescence imaging of deep tissue in freely moving animals. Science (1979) 359, 935–939 (2018).

67. Roe, A. W., Fritsches, K. & Pettigrew, J. D. Optical imaging of functional organization of V1 and V2 in marmoset visual cortex. Anat Rec A Discov Mol Cell Evol Biol **287A**, 1213– 1225 (2005).

68. Tani, T. et al. Sound Frequency Representation in the Auditory Cortex of the Common Marmoset Visualized Using Optical Intrinsic Signal Imaging. eNeuro 5, ENEURO.0078- 18.2018 (2018).

69. Valverde Salzmann, M. F., Bartels, A., Logothetis, N. K. & Schuz, A. Color Blobs in Cortical Areas V1 and V2 of the New World Monkey Callithrix jacchus, Revealed by Non-Differential Optical Imaging. Journal of Neuroscience 32, 7881–7894 (2012).

