## Supplemental Figure 1 for "An Open Access Resource for Marmoset Neuroscientific Apparatus"

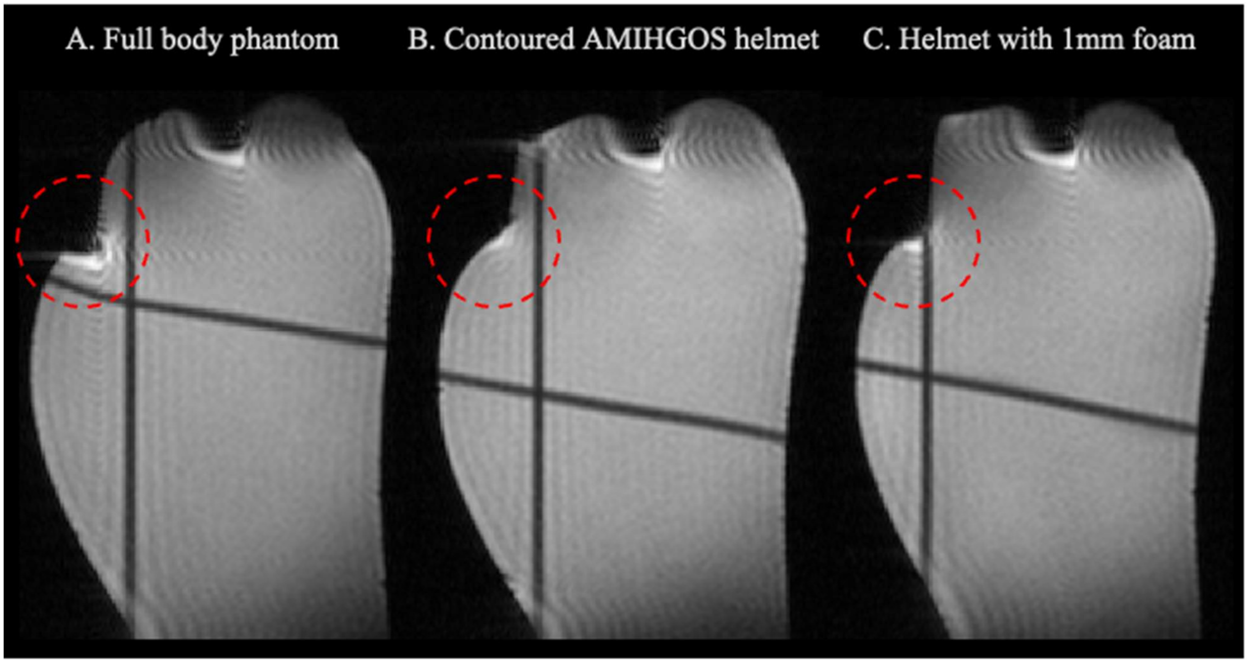


Supplementary Figure 1. (A) Phantom without a helmet, used to visualize existing susceptibility artifacts. (B) 3D printed durable helmet covering the sagittal bump of the phantom corresponding to the brow ridge (red circle). (C) Helmet with an added 1mm foam layer on the inner side, further minimizing susceptibility artifacts. The susceptibility artifact is reduced with the addition of the AMIHGOS-generated helmet, as the interface between the phantom and air is replaced by foam and plastic materials with magnetic susceptibilities closer to that of biological tissue.
